# Inserting small molecules across membrane mixtures: Insight from the potential of mean force

**DOI:** 10.1101/802769

**Authors:** Alessia Centi, Arghya Dutta, Sapun H. Parekh, Tristan Bereau

## Abstract

Small solutes have been shown to alter the lateral organization of cell membranes and reconstituted phospholipid bilayers; however, the mechanisms by which these changes happen are still largely unknown. Traditionally, both experiment and simulation studies have been restricted to testing only a few compounds at a time, failing to identify general molecular descriptors or chemical properties that would allow extrapolating beyond the subset of considered solutes. In this work, we probe the competing energetics of inserting a solute in different membrane environments by means of the potential of mean force. We show that these calculations can be used as a computationally-efficient proxy to establish whether a solute will stabilize or destabilize domain phase separation. Combined with umbrella sampling simulations and coarse-grained molecular dynamics simulations, we are able to screen solutes across a wide range of chemistries and polarities. Our results indicate that, for the system under consideration, preferential partitioning and therefore effectiveness in altering membrane phase separation are strictly linked to the location of insertion in the bilayer (i.e., midplane or interface). Our approach represents a fast and simple tool for obtaining structural and thermodynamic insight into the partitioning of small molecules between lipid domains and its relation to phase separation, ultimately providing a platform for identifying the key determinants of this process.

**SIGNIFICANCE:** In this work we explore the relationship between solute chemistry and the thermodynamics of insertion in a mixed lipid membrane. By combining a coarse-grained resolution and umbrella-sampling simulations we efficiently sample conformational space to study the thermodynamics of phase separation. We demonstrate that measures of the potential of mean force—a computationally-efficient quantity—between different lipid environments can serve as a proxy to predict a compound’s ability to alter the thermodynamics of the lipid membrane. This efficiency allows us to set up a computational screening across many compound chemistries, thereby gaining insight beyond the study of a single or a handful of compounds.

## INTRODUCTION

Many cellular processes, including signal transduction as well as sorting and trafficking of proteins and pathogens, are rooted in the lateral organization of the plasma membrane (1). As purported by the raft concept, ordered and densely packed regions, containing sphingolipids and cholesterol, coexist with regions of loosely arranged phospholipids in biological membranes (2, 3). Artificial membranes, containing adequate amounts of cholesterol, also show formation of similar lipid nanodomains (4, 5). Generally speaking, below the miscibility transition temperature (*T*_mix_), saturated lipids and cholesterol form a phase with a higher degree of order of the hydrocarbon chains, named liquid-ordered (Lo), while unsaturated lipids maintain a more disordered arrangement in the so-called liquid-disordered phase (Ld) (5).

Remarkably, mixing or demixing of the different lipid components can be achieved by incorporating small solute molecules that partition between coexisting domains, thereby shifting phase separation (6, 7). This very intriguing effect, which has been speculated to be linked to the mechanism of action of general anesthetics (6), might have important consequences for cell function, opening a path to the design of new drug-like compounds acting on membrane proteins through lipid domain-mediated effects. Experimentally, the change in lateral organization translates into a shift of the *T*_mix_. This effect has been reported for short-chain alcohols when added to giant plasma membrane vesicles (GPMVs) (6), as well as to ternary giant unilamellar vesicles (GUVs) (7). For both systems the temperature shift is quite significant; however, in opposite direction: short chain alcohols decrease *T*_mix_ in GPMVs while increasing it in GUVs. Interestingly, when the alcohol chain consists of more than 8 carbons the reported shift for GUVs’ *T*_mix_ becomes non-monotonic, making it hard to predict what the effect of a new untested compound will be.

Molecular dynamics (MD) simulations have a long history of complementing experiments when it comes to understanding the intricate details of phase separation in lipid membranes (8–17). The nature of the problem inherently calls for long length-and time-scales. As such, coarse-grained (CG) models that lump together several atoms into one super-particle or bead are particularly suited for this purpose (18). The Martini force field (19, 20) represents a popular choice for studying complex membranes, as demonstrated by a recent review on this topic (21). Using the Martini force fields (19, 20), several computational studies have already investigated the effect of adding specific compounds such as transmembrane peptides (22, 23), amphiphiles (24) as well as small hydrophobic molecules and polymers (25–27) to ternary membranes displaying phase separation. Various possible explanations for the underlying mechanism of domain modulation by small additives have also been proposed. These include: preferential partitioning between Lo and Ld domains (22–24, 26, 27), change in membrane thickness leading to increase or decrease of the hydrophobic mismatch between coexisting phases (24) as well as changes to the line tension at the interface between domains (i.e., linactant mechanism) (25, 28). Taken together, the picture emerging from experimental and computational studies highlights the complexity of the problem, with many possible competing processes occurring simultaneously that could impact domain stabilization or destabilization. To this end, an approach to quickly explore the chemical space at a reduced computational cost while providing simple molecular markers or chemical features that can help predict the bilayer-modifying character of new compounds would be extremely beneficial.

In this work we study, in detail, the thermodynamics of inserting a small solute molecule into a membrane mixture. The potential of mean force (PMF), obtained herein using umbrella sampling (US), provides a robust observable to quantify the stability of the system. Our group previously demonstrated the benefits of calculating PMFs to study the translocation of small molecules in a one-component lipid membrane at high throughput, i.e., across the chemical space of small organic molecules (29–32). PMF measures have otherwise been successfully employed to shed light on the thermodynamic origins of many biologically relevant processes, including protein dimerization (33–36), preferential binding and association of peptides and proteins to different membrane environments (35, 37, 38), as well as to study the selectivity of antimicrobial peptides between bacterial and mammalian-like membranes (39, 40).

We propose to compare different PMFs to study the preferential insertion of a solute between different membrane environments, and its potential ability to shift the Lo-Ld phase equilibrium. In particular we monitor the free energy of inserting the solute in three environments representative of Lo, Ld, and the mixture, and show how they can be indicative of preferential partitioning in large-scale MD simulations. By exploiting the modularity of the Martini force field and the computational efficiency of US, we obtain thermodynamic trends across a wide variety of chemically different compounds. Our results indicate that lipid mixing and demixing originate from a close interplay between preferential partitioning and insertion, where the compounds displaying the strongest effects localize at the bilayer midplane. Ultimately, the combined partitioning and structural information obtained with this approach can lead to a better understanding of the driving forces governing lipid mixing and demixing caused by small molecules.

## METHODS

Coarse-grained (CG) simulations using the Martini force field (20) were carried out with GROMACS 4.6.6 (41) and GROMACS 5.1.2 (42) in combination with PLUMED 2 (43). Two different types of simulations were performed: (*i*) the large-scale reorganization of membrane mixtures under the influence of a small concentration of solute molecules using unbiased MD simulations; and (*ii*) PMF calculations of the insertion of a single small molecule inside a lipid bilayer from umbrella sampling (US). More details about both types of simulations are provided in the following sections.

### Unbiased Molecular Dynamics

MD simulations of a ternary membrane consisting of 1,2-dioleoyl-*sn*-glycero-3-phosphocholine (DPPC), 1,2-dilinoleoyl-*sn*-glycero-3-phosphocholine (DLiPC), and cholesterol (CHOL) at a molar ratio of 7:4.7:5 were carried out using the Martini force field (20, 44). The simulation box contained 612 DPPC, 408 DLiPC, 436 CHOL, and 20732 water molecules, of which 10% were replaced by antifreeze particles. It should be noted that the specific lipid ratio was chosen because it has been shown to reproduce Lo-Ld phase separation (25, 45). The membrane was simulated at different temperatures (289, 295, 305, 310, 315, 325 and 335 K) as well as in the presence of small concentrations (approximately 5 mol%, corresponding to 80 dimer molecules) of solutes at 305 K. The ternary membrane was created using the INSANE building tool (46) while solute molecules were randomly placed in the simulation box. Prior to production runs, all systems were energy minimized, heated up, and equilibrated. Production simulations were then performed in the *NPT* ensemble by keeping the pressure fixed at 1 bar using the Parrinello-Rahman barostat (47) while temperature was controlled using the velocity-rescaling thermostat (48).

### Contact fraction analysis

Following Barnoud et al. (25), we measured the degree of phase separation in each system by calculating the DLiPC-DPPC contact fraction

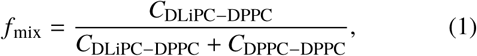

where *C*_*i–j*_ represents the number of contacts between two lipids. Contacts between the phosphate group (PO4 bead) of two lipids are calculated with the GROMACS utility `g_mindist`, using as threshold a distance of 1.1 nm, as reported in previous studies (22, 25).

In a similar fashion we calculated the solute-DLiPC contact fraction, 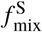, and the cholesterol-DLiPC contact fraction, 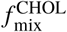, by replacing DPPC in Eq. 1 with the solute or cholesterol, respectively,

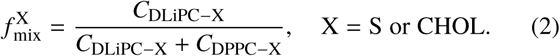

It should be noted that only contacts between phosphate and cholesterol head group (i.e., PO4 and ROH beads) were considered in the calculation of 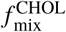, while the entire molecule was considered for the calculation of 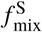. The threshold distance for 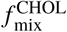 is 1.1 nm, while for 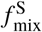. it is reduced to 0.8 nm. Lastly, contact fractions are calculated by averaging over the last 10^7^ *τ* of simulation time (*τ* = 1 ps).

### Umbrella Sampling

US simulations were performed in three different environments: a ternary mixture consisting of DPPC, DLiPC, and CHOL at a molar ratio of 7:4.7:5, referred to as “mix,” and two individual lipid bilayers containing only DPPC or DLiPC, chosen as proxy for the Lo and Ld phases, respectively (20, 44). All membranes were created using the INSANE building tool (46) producing a lamellar system containing 64 lipids per leaflet for DPPC and DLiPC, and 63 randomly distributed lipids per leaflet for the mix system. The lamellar systems were then solvated with Martini water beads. Figure 1-(a) shows the three membrane environments considered. With regards to the solutes, we specifically considered small molecules represented by two connected Martini beads, referred herein as dimers. Rather than focusing on specific compounds, we considered *all* CG dimers by exhaustively enumerating all combinations of neutral Martini beads. This resulted in a total of 105 dimer solutes, covering a wide range of hydrophobicity (29, 30). Hence, a US simulation was constructed for each combination of solute and membrane environment, using as reaction coordinate the distance along the bilayer normal, *z*, between the membrane midplane and the solute molecule. Each simulation consisted of 24 windows spaced every 0.1 nm, with the force constant of the harmonic restraint set to 239 kcal mol^−1^ nm^−2^. To improve sampling, two solute molecules were placed in the simulation box at sufficient distance of one another (49). Prior to production runs, all systems were energy minimized, heated up, and equilibrated. We ran production simulations in the *NPT* ensemble for up to 4 · 10^5^ *τ* at a pressure *P* = 1 bar using the Parrinello-Rahman barostat (47) and a coupling constant *τ*_*P*_ = 12*τ*, while temperature was kept constant at *T* = 300 K using the velocity-rescaling thermostat (48) and a coupling constant *τ*_*T*_ = *τ*.

**Figure 1:**
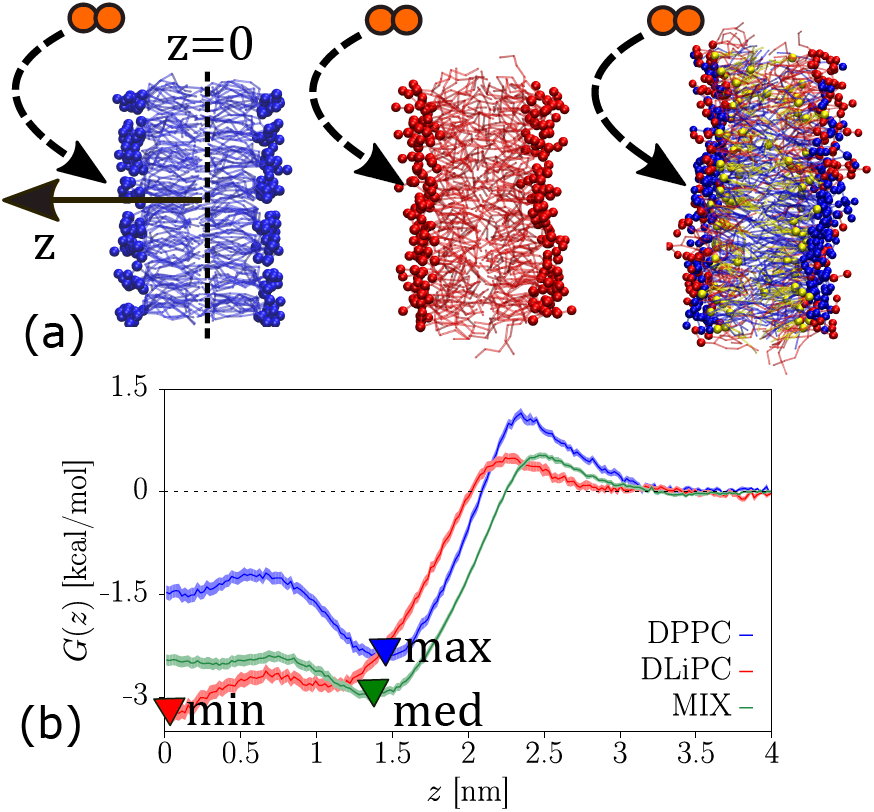
(a) Schematic representation of the protocol employed to perform free-energy calculations. Martini dimers are inserted in three different lipid environments: pure DPPC and DLiPC patches and ternary DPPC:DLiPC:CHOL membrane. Colour code is: DPPC, blue; DLiPC, red and CHOL, yellow. Water is omitted for clarity. (b) An example of PMFs obtained for the dimer C3-N0 in each lipid environment considered (DPPC, blue; DLiPC, red and ternary membrane, green). The three PMFs are labelled as “min,” “med,” and “max” according to the value of their respective transfer free energy from water to the membrane,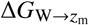.

We relied on the lipid mixture as a proxy for the mixed-membrane phase. This is complicated by the ternary system that naturally demixes at our temperature of interest (see Figure S2-(c)). To ensure sampling of the state point of interest, we controlled the mixing of the membrane via the ratio of contacts between DLiPC and DPPC lipids, i.e., contact fraction *f*_mix_. We applied a harmonic restraint with spring constant *K* = 1195 kcal mol^−1^ nm^−2^, as implemented in PLUMED (43).

PMF profiles were estimated using the weighted histogram method (WHAM) (50) and the relative errors via bootstrapping analysis (51) as implemented in GROMACS (52).

### Umbrella sampling at different cholesterol concentrations

Following the same protocol described in the previous section, we performed US simulations on ternary membranes containing DPPC, DLiPC, and a variable cholesterol concentration. Specifically, we used the same membrane compositions employed by Pantelopulos and Straub to study the effect of cholesterol concentration on lipid membrane phase behavior (53). This corresponds to ternary membranes having an equal ratio of DPPC and DLiPC lipids, while the cholesterol concentration is fixed at 0, 3, 7, 13, 22 and 30 mol% respectively. Each system was prepared for production runs following the same steps described in the previous section. US simulations were performed on a subset of Martini dimers to produce PMF profiles in the mix environment at a variable cholesterol concentration.

### Free-energy calculations

It is first useful to consider the depth at which the solute will preferentially insert, *z*_m_, which corresponds to the minimal value of the PMF, *G*(*z*_m_) = min_*z*_ *G*(*z*). To determine the relative stability of the compound across different membrane environments, we calculate the transfer free energy between water and membrane, 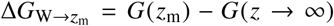, in each system. Subsequently, we identify, for each dimer, which of the three lipid environments produce the largest change in 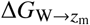, i.e., the lipid environment where the solute will most favourably insert, and denote this as “min.” Similarly we identify the system where solute insertion will be the least favourable (i.e., smallest value of 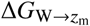), which we denoted as “max,” and the system displaying an intermediate value of 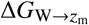 which we refer to as “med.” Figure 1-(b) provides an example of this procedure in the case of the dimer C3-N0. Hence, we calculate the difference in transfer free energies, ΔΔ*G*, between the two environments where each solute inserts more favorably (min and med) to study the possible competition between them (the max environment is therefore discarded). For simplicity of discussion, we attribute a positive sign to ΔΔ*G* if the environment displaying the largest 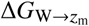 is the ternary system so that

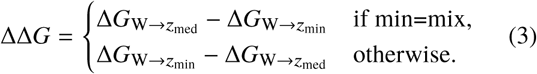

We provide some examples of input data for unbiased MD and US in a repository (54).

## RESULTS

### Mixing and demixing effects induced by small molecules

We carried out MD simulations of the ternary membrane DPPC:DLiPC:CHOL in the ratio 7:4.7:5 and without solute at different temperatures and measured the contact fraction, *f*_mix_. Figure S1 shows the change in contact fraction as a function of temperature. As already reported by Barnoud et al., at low temperature the system appears phase separated (small contact fraction, 0.204 < *f*_mix_ < 0.306), with domains enriched in DPPC and cholesterol (Lo phase) coexisting with domains mostly consisting of DLiPC (Ld phase) (25, 45). Because of the periodic boundary conditions used in our simulations, at the lowest temperatures lipid domains arrange into stripes instead of experimentally-observed patches (see Figure S2 -(a) and (b)). By increasing temperature, we observe enhanced mixing (larger values of contact fraction, 0.343 < *f*_mix_ < 0.471) and a reduction of the stripe-like domains (see Figure S2-(c) to (g)). Since the contact fraction depends on the system size, we cannot directly compare our results with those obtained by Barnoud et al.. Nevertheless, we recover similar trends for *f*_mix_ as a function of temperature. At this point it should be noted that in the remainder of this section we will focus on one specific temperature, 305 K, to evaluate whether phase separation in the ternary membrane is affected by addition of small molecules. Hence, the bilayer at 305 K in absence of any solute effectively represents our reference system and its contact fraction (i.e., *f*_mix_ ≈ 0.31, see Table S1) will be used as a measure to quantify demixing (i.e., *f*_mix_ < 0.31) or mixing (i.e., *f*_mix_ > 0.31) induced by small molecules.

We simulate the ternary membrane in the presence of small solutes, by adding to the system a finite concentration, about 5 mol%, of three different Martini dimers. We chose compounds covering different types of chemistry: one hydrophobic compound (i.e., C1-C1), a compound with intermediate polarity (i.e., C4-C4) and one amphiphilic compound (i.e., C1-Nd). After running each system for 3 · 10^7^ *τ*, we calculate its *f*_mix_ as well as the solute-DLiPC contact fraction, 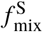, and the cholesterol-DLiPC contact fraction, 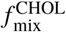.

A summary of the results is provided in Table 1. For each of the solute tested we observe different types of behavior: C1-C1 favors mixing of the ternary membrane (*f*_mix_ = 0.351 ± 0.009), C4-C4 stabilizes lipid domains (*f*_mix_ = 0.271 ± 0.006), while C1-Nd has no significant effect on the phase separation (*f*_mix_ = 0.297 ± 0.005).

**Table 1:** Different contact fractions for Martini dimers C1-C1, C4-C4 and C1-Nd. Δ*G*_Ol→W_ is measured in kcal/mol (55). The contact fraction for the ternary system in the absence of any solute at 305 K is *f*_mix_ ≈ 0.31. Full list of error bars are reported in Table S2.

| Dimer | $\Delta G_{Ol \rightarrow W}$ | $f_{mix}$ | $f_{mix}^S$ | $f_{mix}^{CHOL}$ |
| --- | --- | --- | --- | --- |
| C1-C1 | 6.8 | $0.351 \pm 0.009$ | 0.306 | 0.218 |
| C4-C4 | 4.8 | $0.271 \pm 0.006$ | 0.689 | 0.204 |
| C1-Nd | 4.0 | $0.297 \pm 0.005$ | 0.458 | 0.214 |

With regard to the mixing effect of C1-C1, this result is in line with what was already observed by Barnoud et al., who report an increase in lipid mixing for the DPPC/DLiPC/CHOL system in the presence of aliphatic solutes, amongst which octane, modeled in Martini as a C1-C1 dimer (25). The DLiPC-solute contact fraction (see 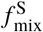 in Eq. 2) indicates which of the two lipids each solute establishes most contacts with, where 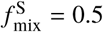 indicates equal contacts with DliPC and DPPC lipids. Hence, by looking at this quantity for C1-C1 dimers 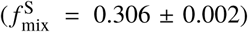 we conclude that they mainly reside in the vicinity of DPPC. As such, C1-C1 preferentially partitions with DPPC, but ultimately leads to mixing, indicating a destabilization effect upon partitioning with the Lo phase. The disruptive effect on phase separation induced by C1-C1 dimers is also clear by looking at Figure 2-(a), where small fragmented DLiPC domains are visible and from the Figure 2-(b), where the density distribution along the *x* direction shows peak shape distortion. Here we also notice that DPPC and DLiPC domains appear slightly anti-registered, and as a result the solute and cholesterol distributions are rather broad.

**Figure 2:**
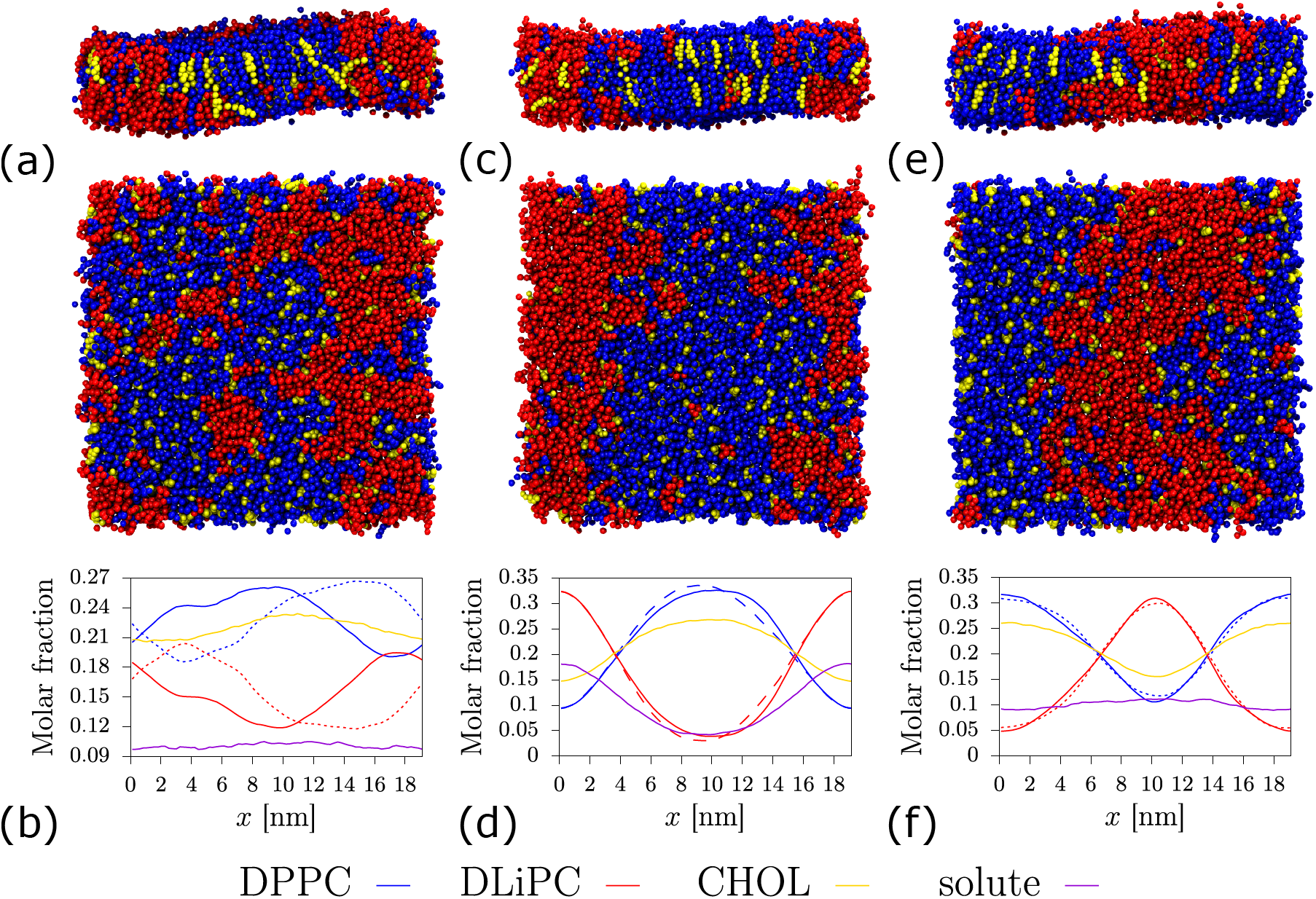
Phase separation in ternary lipid membranes in the presence of small concentrations of dimers: (a) and (b) C1-C1; (c) and (d) C4-C4; (e) and (f) C1-Nd. Snapshots are taken from simulations at 2.8 · 10^7^ *τ* showing the side and top view of the membrane. (b), (d) and (f): density profiles in the direction of phase separation expressed as molar fractions and averaged over the last 10^7^ *τ* of simulation time. Continuous and dashed lines are used to distinguish between the two leaflets. DPPC: blue; DLiPC: red; CHOL: yellow; and solute: purple.

On the contrary, C4-C4 dimers preferentially partition with DLiPC lipids 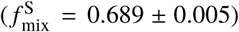 and cause *f*_mix_ to decrease in comparison to the pure DPPC/DLiPC/CHOL membrane, an indication of more stable phase separation. Lo and Ld domains are clearly visible in Figure 2-(c) and the overall structure of the membrane appears rather ordered, with clear distinguishable lipids and cholesterol peaks and accumulation of the solute in the DLiPC phase (see Figure 2-(d)).

Lastly, C1-Nd dimers display approximately equal partitioning between DPPC and DLiPC 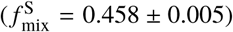, as evident by the solute distribution in Figure 2-(f), and produce effectively no change to the Lo/Ld phase separation (see Figure 2-(d)).

The effects observed for the three above-mentioned dimers indicate that a relationship exists between preferential partitioning (or lack thereof) in different membrane environments and the solute mixing or demixing character. Specifically, we note that a dimer that localizes near DPPC lipids (i.e., the main component of Lo phase) enhances lipid mixing in a phase separated ternary membrane; on the contrary, when the dimer localizes near DLiPC lipids (i.e., the main component of Ld phase) we observe domain stabilization in the same ternary membrane, and lastly no significant modification to phase separation is observed for dimers that partition approximately equally between DPPC and DLiPC lipids. We also notice that the DLiPC-CHOL contact fraction, 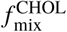, does not change significantly with respect to the solute-free system (see Table 1 and Table S1). We conclude that the transfer of cholesterol from Lo to Ld phase is rather moderate here. In the next section we investigate this aspect further by evaluating the thermodynamics of insertion of small molecules in different lipid membranes by means of US simulations.

### Predicting phase separation from several potentials of mean force

The protocol described in the previous section aims at establishing whether a solute stabilizes or destabilizes membrane phase separation. Based on large-scale MD simulations, it is unfortunately computationally demanding and becomes impractical when screening larger numbers of molecules. Here we seek a computationally efficient and insightful proxy to this protocol. We argue that the structural and thermodynamic information contained in PMFs can be leveraged to estimate the *relative* stability of a compound between different environments: the membrane mixture and one-component domains (i.e., DPPC for Lo and DLiPC for Ld). Because these different PMFs are calibrated against a common environment (i.e., bulk water), relative transfer free energies can be used as a proxy for the preferential stability of a compound in an environment. We thereby hypothesize a link between the maximum transfer free energy from water to one of the three membrane environments and the propensity to drive membrane phase separation.

We studied 16 Martini dimers using both approaches: (*i*) a large-scale MD simulation and (*ii*) PMF calculations in the three different membrane environments. We chose dimers with different levels of hydrophobicity, but avoid strongly-polar dimers (e.g., P-P type), which do not favorably insert in the membrane. Hence, for each dimer we calculate the difference, ΔΔ*G*, between two most favorable transfer free energies, defined as “min” and “med” environments, as described in Eq. 3. Additionally, the mixing or demixing character of each solute is characterized by the DLiPC-DPPC contact fraction, *f*_mix_, measured from the unbiased MD simulations (see Eq. 1).

We find that the information contained in the PMFs indeed correlates with the large-scale MD simulations. We find a linear correlation between the free-energy difference measured between competing environments and the contact fraction, as shown in Figure 3-(a). In particular we observe that dimers that prefer the mixed environment (i.e., positive ΔΔ*G*) yield a larger contact fraction, while dimers favoring a one-component membrane (i.e., negative ΔΔ*G*) lead to a decrease in the contact fraction. Furthermore, plotting the change in ΔΔ*G* against the solute-DLiPC contact fraction 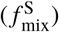 in Figure 3-(b) reveals that, in agreement with our findings described in Mixing and demixing effects induced by small molecules, dimers that produce mixing (i.e., positive ΔΔ*G*) localize near DPPC 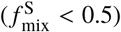 while dimers enhancing demixing (i.e., negative ΔΔ*G*) localize near DLiPC 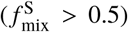 Interestingly, all dimers that affect phase separation preferably localize at the membrane midplane, see purple points in Figure 3. Here each dimer has been colored according to their *z*_min_, i.e., the value of *z* in the environment displaying the largest ΔΔ*G*. We notice that indeed for ΔΔ*G* values between approximately – 0.25 and 0.25 kcal/mol, which is indicative of rather moderate preference for one lipid environment, *z*_min_ > 1.2 nm and the dimer localizes at the interface, while dimers that insert into the midplane region (*z*_min_ < 0.5 nm) display larger values of ΔΔ*G*. This results is consistent with the fact that DPPC and DLiPC only differ in their tails. In other words, a dimer that preferentially resides at the interface will not be able to distinguish between the two lipid species, as shown by the small differences in ΔΔ*G* between min and med environments and 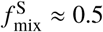.

**Figure 3:**
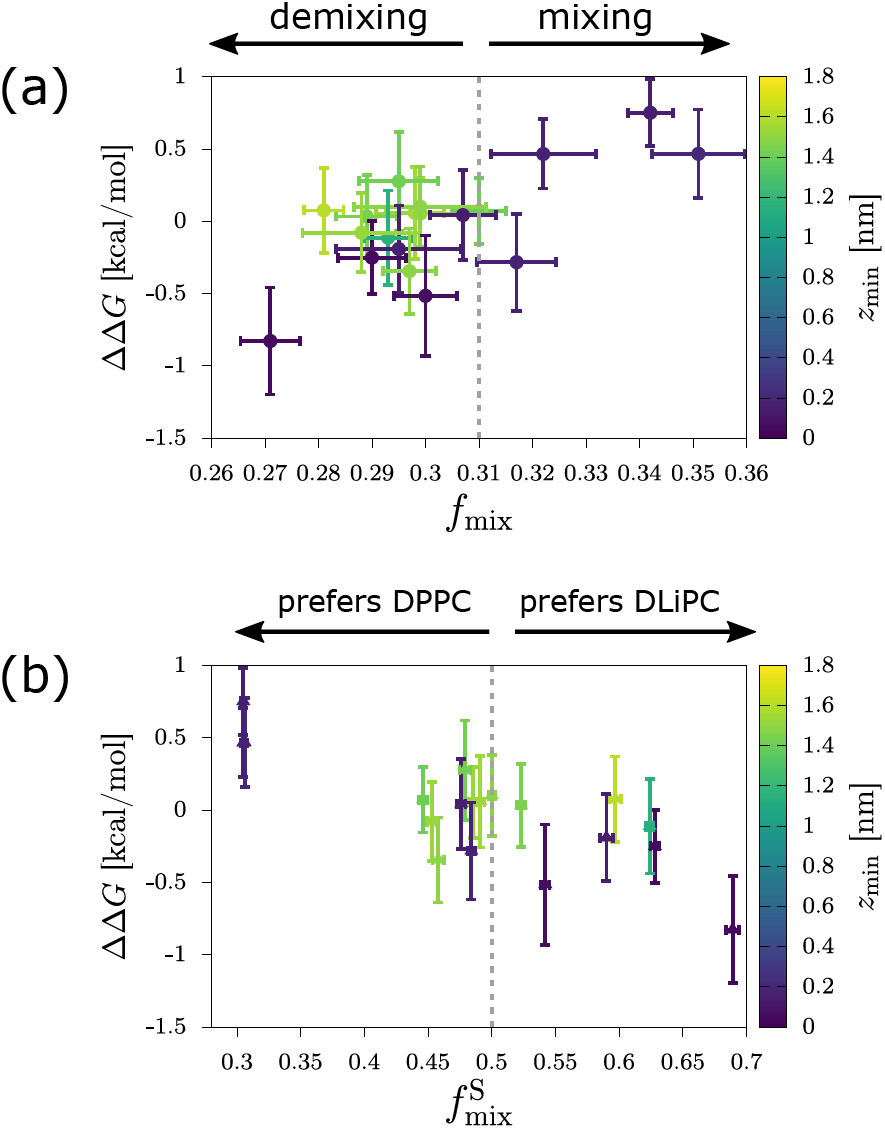
ΔΔ*G* difference between min and med environments as a function of *f*_mix_ (a) and 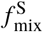 (b) for selected dimers. Solutes are colored according to their respective *z*_min_ location, where *z* = 0 corresponds to the membrane midplane. The dashed grey lines indicate (a) the contact fraction for the system at 305 K in absence of solutes, *f*_mix_ = 0.31, and (b) the solute-DLiPC contact fraction for equal partitioning of the solute, 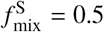.

### High-throughput search for phase-modifying solutes

We have shown in the previous section that the strength of phase separation, expressed by *f*_mix_, correlates well with the dimer’s relative partitioning between different lipid environments, quantified by the transfer free energy, ΔΔ*G*. As such, using PMFs in the three lipid environments representative of Lo-Ld equilibrium offers two advantages: (*i*) identify the preferred lipid environment and (*ii*) estimate the dimer-induced effects on phase separation, both at a reduce computational cost (29–31). Additionally, the PMF profile contains spatial information about the insertion, which we have observed to also play a role in determining the bilayer-modifying character of the solute.

We hereby extend our PMF analysis to predict the strength of phase separation to the larger dataset of all neutral Martini dimers—105 compounds in total—to further understand the origin of mixing and demixing effects induced by small molecules. Hence, for each dimer-bilayer system we measure the transfer free energy ΔΔ*G*, leading to the results displayed in Figure 4.

**Figure 4:**
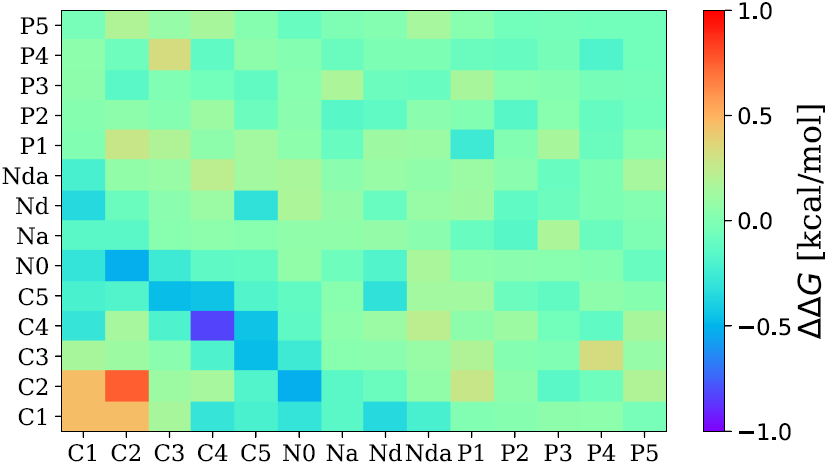
Two-dimensional matrix showing the ΔΔ*G* value across the three lipid environments considered (DPPC, DLiPC and mixed ternary membrane) for all neutral Martini dimers. Horizontal and vertical axes show the bead type combination of each dimer. The grid is symmetrical across the diagonal. The sign of ΔΔ*G* is positive when the lipid environment displaying the largest 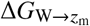 is the ternary system (see Eq. 3).

Two main areas of interest can be identified: (*i*) the orange-red square comprising the most hydrophobic dimers (i.e., C1-C1, C1-C2 and C2-C2) at the bottom left corner, and (*ii*) the blue diagonal stripe comprising dimers with intermediate polarity (approximately 4.2 kcal/mol, i.e., C2-N0, C3-C5, C4-C5 and C4-C4). According to our sign convention for ΔΔ*G*, the dimers belonging to the first group have a preference for the mixed membrane (i.e., positive ΔΔ*G*) and increases lipid mixing, while the second group of dimers favors the pure lipid systems (i.e., negative ΔΔ*G*) causing membrane demixing. Besides these two regions, the remainder of the matrix displays values of ΔΔ*G* close to zero, indicating approximately equal preference for the two most favorable environments and no significant alteration to phase separation. It is worth pointing out that, for each dimer combination in Figure 4, Δ*G*_O1→W_ decreases moving along the lower-left to upper-right diagonal, as shown in Figure S3. This indicates that, as the hydrophobicity content decreases, the dimers gradually shift their mixing-to-demixing character as well as their location of insertion from midplane to interface (cf. Figure S3, Figure 4 and Figure S4 -(b)).

To assess the robustness of the results shown in Figure 4, we compared the transfer free energies for dimers made of two C-type beads in membranes prepared at different cholesterol concentrations. The small concentration changes we apply are such that we do not expect significant changes in the observed trends (see Umbrella sampling at different cholesterol concentrations for more details). The chosen lipid-to-cholesterol ratios match the study of Pantelopulos and Straub, who have recently identified different regimes of phase separation for the DPPC/DLiPC/Chol system modeled using the Martini force field (53). Specifically, they observe a stabilization of the Ld phase at low-cholesterol concentration (0-3 mol%); the onset of phase separation at 7 mol% of cholesterol, with coexisting Lo-Ld domains persisting up to 42 mol%. Finally, at very high cholesterol concentrations, significant anti-registration is observed and Ld domains coexist with a newly identified “cholesterolic” gel phase (53). We are interested in the Lo-Ld regimes of phase separation, accordingly we only measure PMFs for systems with cholesterol concentrations ranging from 0 to 30 mol%. Figure S7 shows that small changes in cholesterol composition do not significantly alter the water-membrane transfer free energy. Additionally, the two dimers that previously displayed the strongest tendency to alter phase separation (i.e., C2-C2 favouring the mixed state and C4-C4 favouring the pure patch), persist in displaying remarkable characteristics at different lipid-to-cholesterol ratios. This effect is, however, significantly weaker for the three lowest cholesterol concentrations at which the system does not yet show stable Lo-Ld domains (see Figure S8). Indeed, the system at 30 mol% cholesterol, the closest in terms of composition to our original system, shows good agreement with the ΔΔ*G* values in Figure 4.

More insight into the relationship between dimer polarity and alteration characteristics of the phase separation can be obtained by probing the preferential location of partitioning, *z*_min_. Two aspects become apparent when we consider the dimers with the largest ΔΔ*G* values (positive or negative): (*i*) they all localize close to the midplane (*z*_min_ < 0.5 nm) in their preferred lipid environment (see Figure 5-(a)) and (*ii*) more dimers reside at the interface (*z*_min_ > 1.5 nm) in the med environment compared to the min environment (cf. Figure 5-(a) and (b)). Unsurprisingly, the most hydrophobic dimers of the dataset (e.g., C-C types) insert close to the bilayer midplane (see Figure S5). However, individual PMF profiles reveal that moderately amphiphilic solutes (4.1 ≲ Δ*G*_Ol→W_ ≲ 4.8 kcal/mol) *change* their preferred location inside the bilayer, depending on the type of lipid environment. As the ratio of unsaturated lipids in the system increases (i.e., DPPC →mix → DLiPC), moderately amphiphilic compounds move from the membrane-water interface to the bilayer midplane (Figure S6). Interestingly, this change in depth of insertion is observed for all dimers previously identified as phase separating (i.e., blue-diagonal stripe in Figure 4). This suggests that domain stabilization depends not only on preferential partitioning but also on its localization. As a general trend, the majority of dimers we probed stabilize at the interface (see Figure S4). This includes amphiphilic compounds consisting of one C-type connected to a N-or P-type bead, as hydrophobic and hydrophilic parts, respectively, as well as compounds with intermediate polarities (N-N type), which do not fully insert in the lipid membrane. The most polar dimers of the dataset (type P-P and N-P), on the other hand, do not favorably reside at the interface (*z*_min_ > 2.5 nm). We note that the localization parameter, *z*_min_, does not significantly change with cholesterol concentration (see Figure S9).

**Figure 5:**
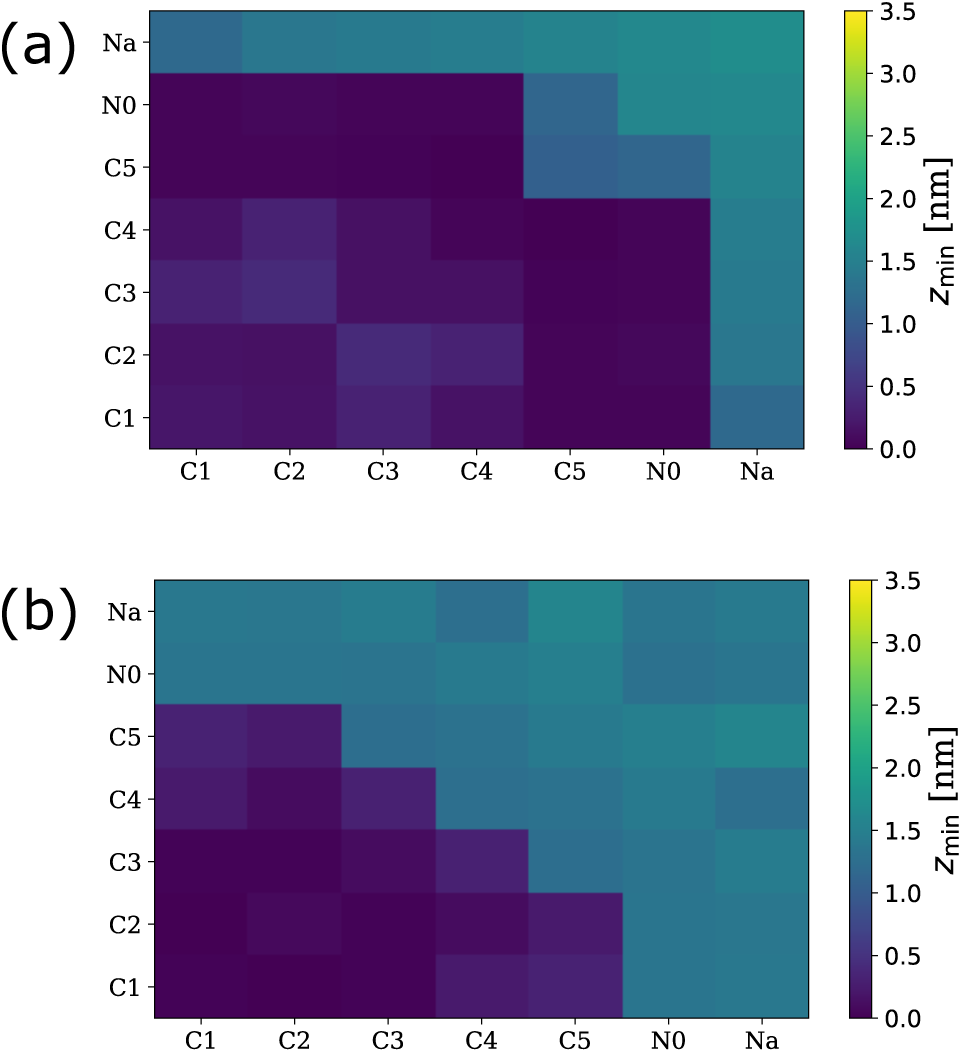
Two dimensional matrices showing *z*_min_ location of a subset of dimers in the two most favoured environments: minimum (min) environment (a) and median (med) environment (b). Horizontal and vertical axes show the bead type combination of each dimer. The grids are symmetrical across the diagonal. The full matrices are provided in Figure S4.

### Mean-field model

In this section we try to rationalize the dimers’ behavior, as well as other observations from our simulations, by using a recent analytical mean-field study. Using a Flory-Huggins type of mean-field free energy, Allender and Schick studied the change in the miscibility transition temperature (Δ*T*_mix_) of a lipid bilayer composed of an unsaturated (A) and a saturated (B) lipid upon addition of a solute (S) (56). The purpose of using a simplified mean-field model is to find out *how* the thermodynamic driving forces (namely, direct and excluded-volume interactions between the solute and the lipids) affect the solute’s ability to phase separate or mix the bilayer. To this end, they calculated Δ*T*_mix_ ^1^ long the critical line on the surface of coexistence separating the mixed and demixed phases and found

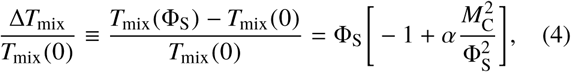

to first order in the solute volume fraction, Φ_S_ (see SI section Mean-Field Theory for details). In the above equation *T*_mix_ (Φ_S_) and *T*_mix_ (0) are the critical mixing temperatures in the presence and absence of solute molecules, respectively; *α* = *k*_B_*T*_mix_ (0)/(2*N*_S_*V*_AB_) is a parameter that depends on the pairwise interaction energies, *V*_AB_, between A and B type monomers in the mixture; *k*_B_ is the Boltzmann constant and *N*_S_ is the number of monomers in a solute molecule. *M*_C_ is proportional to the critical partitioning of the solute in the Ld and Lo phases (see Eq. (6)). Allender and Schick found it to be determined by both excluded-volume (*δv*) and direct (*δr*) interactions

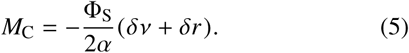

In the above equation, *δv* represents the excluded-volume interactions of the lipid components; it depends only on the number of monomers per lipid chain (*N*_A_ for lipid A and *N*_B_ for lipid B) and it is given by 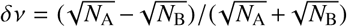. *δr* represents the direct interaction between the lipid and solute and it is found to be *δr* = (*V*_AS_ − *V*_BS_)/*V*_AB_. From Eq. (4) we note that the change in the miscibility transition temperature upon adding solutes is determined by the competition between two terms: a dilution effect and the preferential partitioning of the solute in either phase. While dilution simply arises from the reduced interactions between lipid molecules by the presence of the solute, the preferential partitioning stems from a non-zero value of *M*_C_. When the solute partitions equally into either of the phases (i.e., *M*_C_ = 0), Eq. (4) predicts the largest decrease in the miscibility transition temperature, making Δ*T*_mix_ large and negative. Solutes which prefer one lipid environment, however, partition between the mixed and demixed phases according to their interaction energy and therefore have a finite value of *M*_C_ (see Eq. (5)). Solutes that partitions unequally, but only weakly so, have lower values of *M*_C_ and, from Eq. (4), if 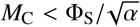, then Δ*T*_mix_ becomes negative and induces mixing. Conversely, for strongly partitioning solutes, specifically, if |*M*_C_| is larger than 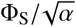, Δ*T*_mix_ becomes positive and induces demixing. Thus even without changing a lipid bilayer’s temperature, it is possible to induce mixing or demixing by only adding a solute to it. Also, according to the mean-field model, the solute’s ability to initiate this transition depends on how preferentially it partitions and that, in turn, is decided by the interplay of the excluded-volume and direct interactions between the solute and the lipids as shown in Eq. (5).

We now relate the mean-field model to our computer simulations, associating lipids A and B to DLiPC and DPPC, respectively. To qualitatively estimate the excluded-volume interactions for our system we first observe that, due to its tail’s higher degree of unsaturation, DLiPC has a kink in its tail and, consequently, has a larger effective volume in the mixture (hydrophobic volume) than that of DPPC. In their paper, Allender and Schick noted that the number of monomers in a lipid chain approximately represents its hydrophobic volume. The hydrophobic volume mismatch between DLiPC and DPPC can then be accounted for by taking *N*_DLiPC_ > *N*_DPPC_ (i.e., *N*_A_ > *N*_B_). This implies a positive excluded-volume term δ*v*. Also, as we are not changing the lipid mixture in our simulations, the excluded-volume contribution to *M*_C_ for different dimers remains constant within the mean-field formalism (see the definition of δ*v* above), while the direct interaction contribution, *δr*, changes as it does depend on the chemistry of the compound. We identify two distinct scenarios, corresponding to mixing and phase-separating solutes. We note that *M*_C_ can be expressed as

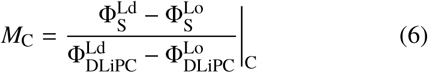

where 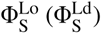 and 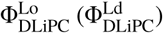 are the *critical* volume fractions of the solute and DLiPC lipids, respectively, in the Lo (Ld) phase near the miscibility transition temperature. Each of the volume fractions in Eq. (6) can be estimated from our simulations (see Eqs. (S5-S8) for the definitions), leading to an estimate of *M*_C_. We can then calculate Δ*T*_mix_ for each dimer using the obtained value of *M*_C_ in Eq. (4). If the mean-field model’s results are consistent with our simulations, then the phase-separating dimers we found in our simulations (classified so based on t heir *f*_mix_ value), will have a positive Δ*T*_mix_ and the dimers promoting mixing will have a negative Δ*T*_mix_.

We compare the mean-field p redictions w ith o ur simulations in Figure 6. We probe the change in miscibility transition temperature, Δ*T*_mix_, as a function of the system’s phase separation, *f*_mix_. To this end we relate *f*_mix_ to *M*_C_ (see Figure S10). Now, in addition to *M*_C_, we also need to know Φ_S_ and *α* to calculate Δ*T*_mix_ using Eq. (4). First, we estimate Φ_S_ = *n*_S_ *N*_S_/(*n*_A_ *N*_A_ + *n*_B_ *N*_B_ + *n*_S_ *N*_S_) = 0.0192. We then note that *α* is adjustable and is given by *α* = *k*_B_*T*_mix_ (0)/(2*N*_S_*V*_AB_).

**Figure 6:**
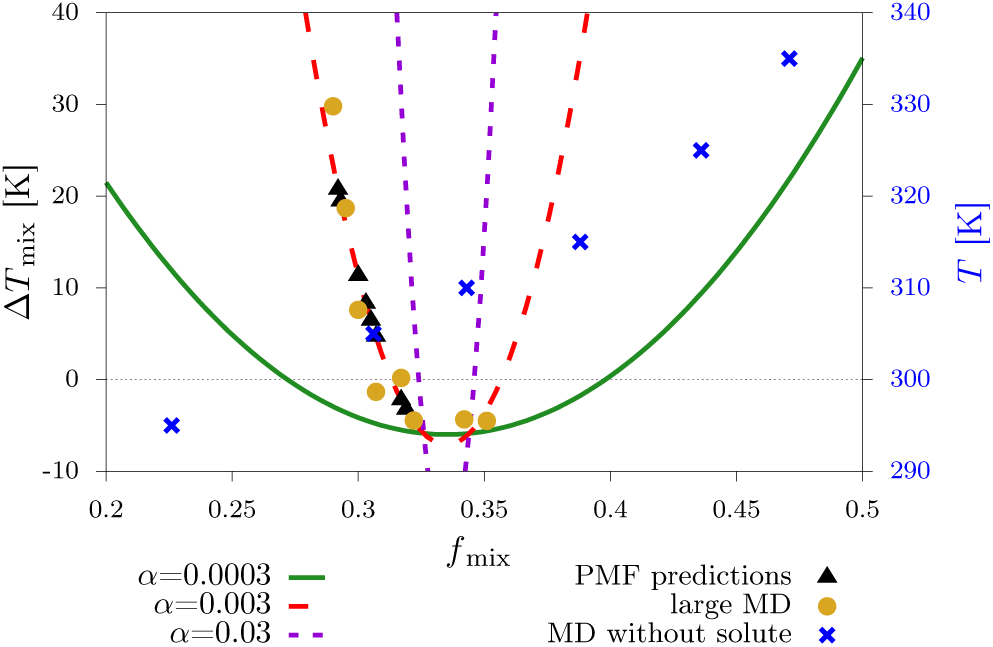
Comparison between simulations and mean-field theory. The change to the miscibility transition temperature in the presence of a finite solute concentration, Δ*T*_mix_, is plotted against the system phase separation, *f*_mix_. Each curve corresponds to a different fit of the adjustable parameter *α*. The yellow circles and black triangles are obtained from large-scale MD simulations and PMF predictions, respectively. The blue crosses, linked to the *T* axis shown on the right, represent MD simulations in the absence of solute at different temperatures (see Table S1and Figure S1).

Assuming a reasonable Δ*T*_mix_ will be about 10 − 30K, we set *α* = 0.003 (red curve). This also fixes the interaction energy *V*_AB_ ≈ 80*k*_B_*T*. For comparison, while *α* = 0.03 results in a lower, more plausible value of *V*_AB_ ≈ 8*k*_B_*T*, it overestimates Δ*T*_mix_ for phase separating dimers (violet curve). A lower *α* = 0.0003 predicts a low Δ*T*_mix_ which incorrectly classifies some phase-separating dimers as mixing ones (green curve), and moreover, strongly overestimates *V*_AB_ ≈ 800*k*_B_*T*. In Figure 6, the yellow circles report on large-scale MD simulations of various Martini dimers. The black triangles complement this subset for compounds where we did not run large-scale simulations, but instead estimate *f*_mix_ from ΔΔ*G* (Figure 3). Focusing on specific compounds and assuming *T*_mix_ (0) ≈ 305 K, the maximum positive shift in the mixing temperature happens for C4-C4, *T*_mix_ (Φ_S_) ≈ 338 K, while the maximum negative shift in the mixing temperature happens for C1-C1, *T*_mix_ (Φ_S_) ≈ 300 K. For reference we have also shown *f*_mix_ values and corresponding bilayer temperatures in the *absence* of solute (blue crosses). We note that changing the system’s temperature leads to a range 0.2 < *f*_mix_ < 0.5 that is much larger than what we have observed by introducing solutes (≈ 0.25 – 0.35). While we find that *M*_C_ decreases linearly with respect to *f*_mix_ (see Figure S10), extrapolating this behavior would lead to Δ*T*_mix_ > 0, implying phase-separation, while an increase in *f*_mix_ corresponds to increased mixing. Further studies in that regime would be useful to more broadly check the mean-field predictions.

The mechanisms obtained from the computer simulations can now be compared to the mean-field theory. First, all simulated compounds that affected the mixing or demixing behavior preferentially inserted at the bilayer midplane—a likely consequence of the composition we probe. This incidentally better fits within the scope of the mean-field model (i.e., no interfacial effects). In our simulations, dimers that promote phase-separation in the bilayer (C2-N0, C3-C5, C4-C5, and C4-C4) partition to the DLiPC-rich Ld phase (see Table S3). The mean-field model is in line with this observation: dimers displaying direct interactions that dominate over excluded-volume interactions (specifically when *δr* < – δ*v*, see Eq. (5)) yield large positive values of *M*_C_. According to Eq. (6), these compounds prefer the DLiPC-rich Ld phase, leading to Δ*T*_mix_ > 0 (see Eq. (4))—thus inducing phase separation. The other bilayer-altering behavior is observed in both cases: CG dimers that promote mixing in the bilayer (C1-C1, C1-C2, and C2-C2) largely partition to the DPPC-rich Lo phase. Likewise within the mean-field model when solutes display weak direct interactions (specifically when *δr*> – δ*v*): even weak preference for the DLiPC-rich Ld phase will seem negligible compared to the excluded-volume interactions. This yields *M*_C_ ≲ 0, so that the solute indeed partitions into the DPPC-rich Lo phase (see Eq. (6)), leading to mixing: Δ*T*_mix_ < 0 (see Eq. (4)). On the other hand, our CG simulations identify a number of dimers (e.g., C3-C3, C3-C4, C3-N0) that, despite reaching the bilayer midplane, did not alter the thermodynamics of the bilayer in any significant way: we observe ΔΔ*G* ≈ 0 between competing lipid environments. This absence of phase-separating or mixing effect for *almost* equally-partitioning solutes does not seem to agree with the predictions of the mean-field model. Instead, solutes that equally partition are expected to produce the maximum *decrease* in *T*_mix_, due to a pure dilution effect (see Eq. (4)) (56).

## CONCLUSION

We have investigated the effect of small molecules on membrane-phase separation for the system DPPC/DLiPC/CHOL using a combination of coarse-grained simulations, unbiased molecular dynamics, and umbrella sampling simulations. By taking advantage of several potential-of-mean-force calculations, corresponding to the insertion of a solute in distinct lipid environments, we have identified a linear relationship between preferential partitioning and phase separation, quantified by the contact fraction, *f*_mix_. Our results show that the phase-modifying character of certain solutes correlates with the difference in transfer free energies between competing lipid environments and that partitioning to the bilayer midplane (*z*_min_ < 0.5 nm) is crucial to produce any alteration to the phase separation. Specifically, we found that dimers that partition to the midplane of the Ld phase act as domain stabilizers, while dimers that partition to the midplane of the Lo phase enhance lipid mixing, in agreement with previous simulation studies (22, 24, 26, 27). By migrating to the DLiPC midplane, stabilizing compounds can occupy regions of the membrane inherently more disordered and where more space is available for localization, ultimately acting as domain stabilizers. The opposite is true for compounds that increase mixing: they also preferentially localize in the midplane region, but do so in the Lo phase, where they compete with the favorable interactions of DPPC with cholesterol, thereby disrupting the Lo domains’ ordered structure. Furthermore, the non-bilayer-modifying character of interfacial solutes (*z*_min_ > 1.5 nm) can be rationalized by taking into account that DPPC and DLiPC lipid heads are identical, and thus insertion close to the interface does not allow for significant preferential partitioning.

Comparison of our simulation results with a Flory-Huggins type mean-field theory (56) helped us rationalize the change in the miscibility transition temperature introduced by the addition of solutes. We find several regimes where the simulation results match with the mean-field predictions: solutes that preferentially partition in the Ld phase induce demixing, while solutes that moderately prefer the Lo phase induce mixing. However, our simulations also report on dimers that partition approximately equally between competing lipid environments, but do not alter the thermodynamics of the lipid bilayer, while the mean-field model predicts a maximum decrease in the miscibility transition temperature, purely due to dilution. This regime was not observed in our simulations, possibly due to the small-solute concentration used. In this regard, Barnoud et al. have reported an increased tendency to mix (i.e., larger contact fraction values) as the solute concentration is increased, for the same lipid mixture (25).

Ultimately, by extracting chemical features and attributes that are characteristic of a certain mixing or demixing behavior, our high-throughput study provides a simple computational approach for the rapid classification of solute molecules interacting with a ternary lipid membrane, as well as insight into the relevant driving forces.

## AUTHOR CONTRIBUTIONS

TB and SP designed the research. AC and AD carried out all the simulations and analyzed the data. All authors wrote the article.

## ACKNOWLEDGMENTS

We thank Robin Cortes-Huerto, Martin Girard, Roberto Menichetti and Nikita Tretyakov for useful discussions and critical reading of the manuscript. The work was partially supported by BiGmax, the Max Planck Society’s Research Network on Big-Data-Driven Materials-Science. T.B. was supported by the Emmy Noether program of the Deutsche Forschungsgemeinschaft (DFG).

## Footnotes

1 While Allender and Schick defined this quantity as a dimensionless ratio, here we define it as a temperature difference.

