## Supplementary Material for "Inserting small molecules across membrane mixtures: Insight from the potential of mean force"

#### Change in phase separation as a function of temperature in the system DPPC/DLiPC/CHOL in absence of solutes

We have performed MD simulations of the ternary membrane DPPC:DLiPC:CHOL in the ratio 7:4.7:5 and in absence of solutes at different temperatures. To evaluate the change in phase separation the DLiPC-DPPC contact fraction, $f_{\mathrm{mix}}$, is calculated for each system (see Unbiased Molecular Dynamics). Figure S1 shows the change in contact fraction as a function of temperature.

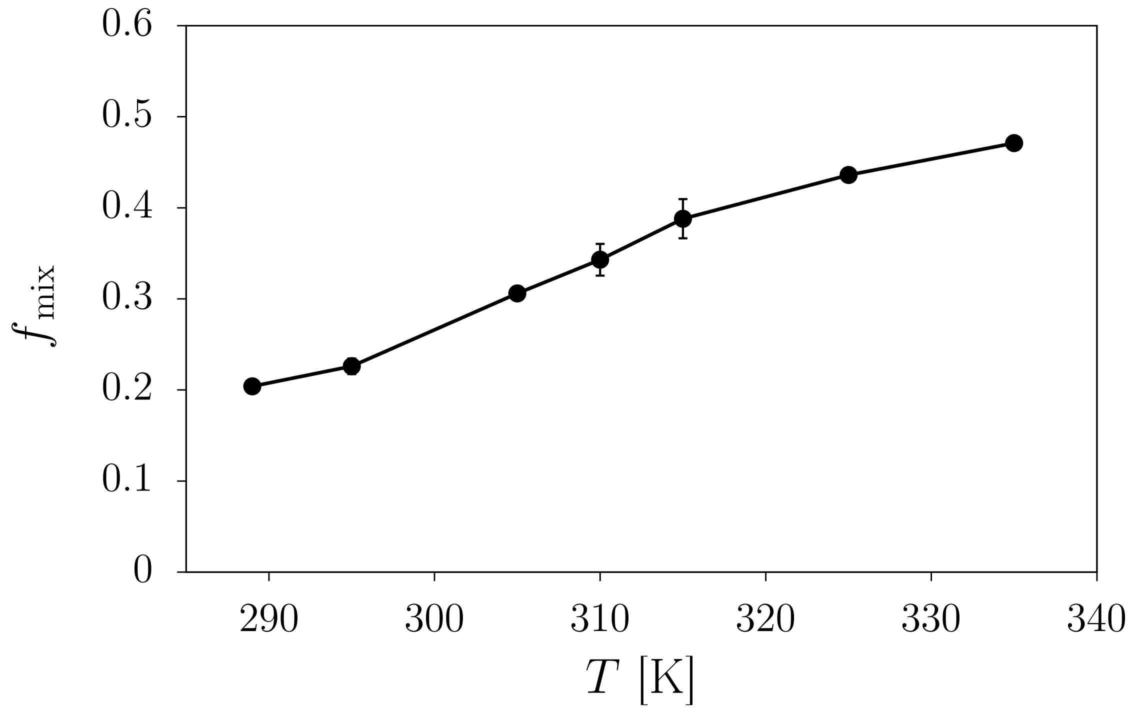


Figure S1: Contact fraction $f_{\mathrm{mix}}$ for the system DPPC:DLiPC:CHOL as a function of temperature as measured in unbiased MD simulations in absence of solutes.

In Table S1 the DLiPC-DPPC contact fraction, $f_{\mathrm{mix}}$, and the DLiPC-CHOL contact fraction, $f_{\mathrm{mix}}^{\mathrm{CHOL}}$, at different temperatures are reported.

Table S1: Contact fraction values for the system DPPC:DLiPC:CHOL obtained form simulations at different temperatures.

| T | $f_{\mathrm{mix}}$ | $f_{\mathrm{mix}}^{\mathrm{CHOL}}$ |
|---|---|---|
| 289 | 0.204 ± 0.0031 | 0.175 ± 0.0020 |
| 295 | 0.226 ± 0.0086 | 0.186 ± 0.0032 |
| 305 | 0.306 ± 0.0060 | 0.210 ± 0.0014 |
| 310 | 0.343 ± 0.0174 | 0.224 ± 0.0024 |
| 315 | 0.388 ± 0.0215 | 0.236 ± 0.0062 |
| 325 | 0.436 ± 0.0024 | 0.259 ± 0.0010 |
| 335 | 0.471 ± 0.0066 | 0.274 ± 0.0024 |

Figure S2 shows the final snapshot for each system.

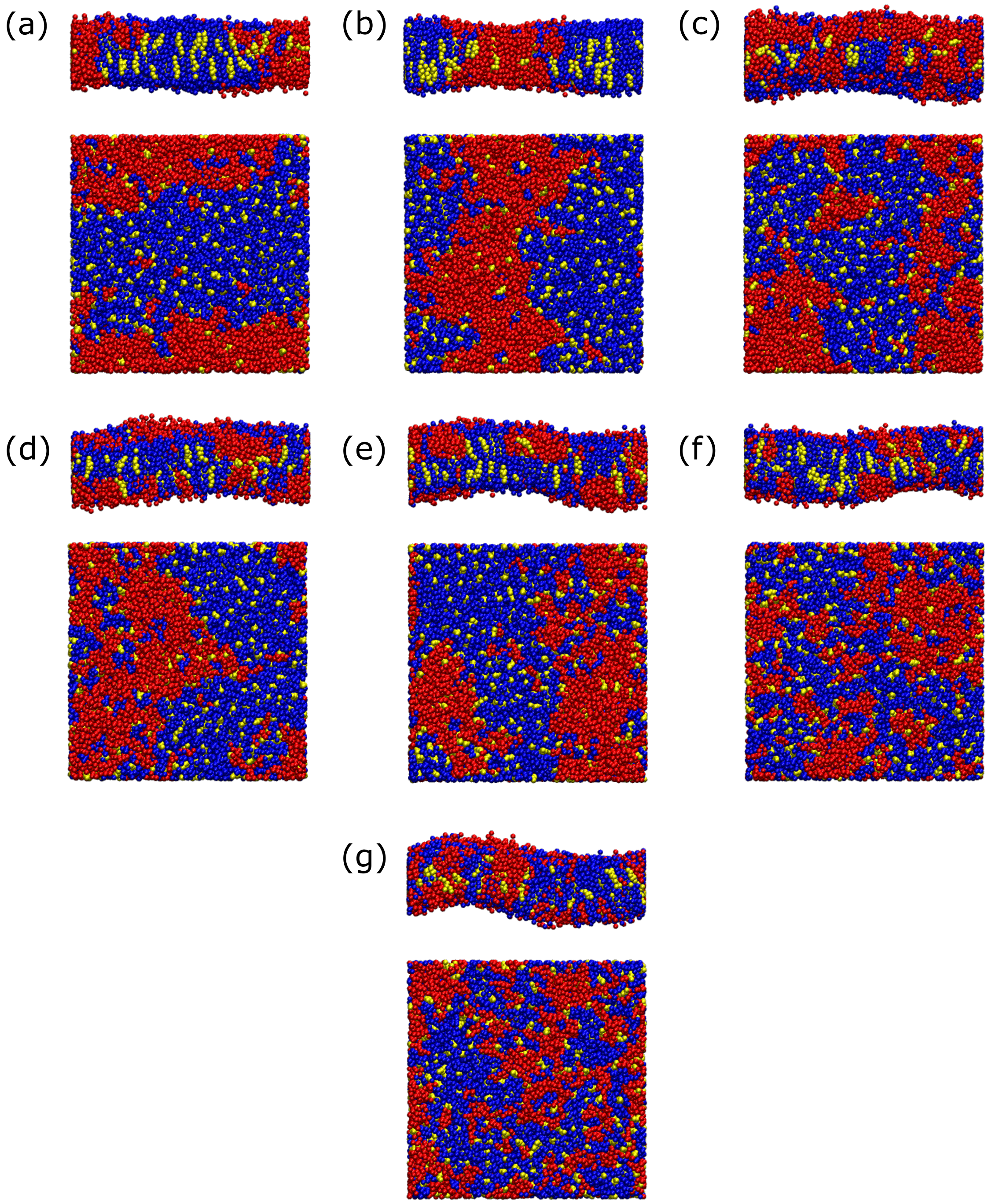


Figure S2: Final snapshots from simulations at (a) 289, (b) 295, (c) 305, (d) 310, (e) 315 and (f) 325 and (g) 335 K for the system DPPC:DLiPC:CHOL, showing the side and top view of the membrane. Colour code is the same as in Figure 1.

### MD Simulations with different dimer types at 305 K

Table S2 shows DLiPC-DPPC contact fraction, $f_{\mathrm{mix}}$, solute-DLiPC contact fraction, $f^{\mathrm{S}}_{\mathrm{mix}}$, and $z_{\mathrm{min}}$ location obtained from simulations with different Martini dimers at 305 K. See Eq. 1 and Eq. 2 for details.

Table S2: Contact fractions, $f_{\mathrm{mix}}$ and $f^{\mathrm{S}}_{\mathrm{mix}}$, and $z_{\mathrm{min}}$ location obtained from simulations with different dimer types at 305 K. For each dimer the octanol/water partition free energy, $\Delta G_{\mathrm{Ol}\rightarrow\mathrm{W}}$, measured in kcal/mol is also is reported [2].

| Dimer | $\Delta G_{\mathrm{Ol}\rightarrow\mathrm{W}}$ | $f_{\mathrm{mix}}$ | $f^{\mathrm{S}}_{\mathrm{mix}}$ | $f^{\mathrm{CHOL}}_{\mathrm{mix}}$ | $z_{\mathrm{min}}$ [nm] |
|---|---|---|---|---|---|
| C1-C1 | 6.8 | 0.351 ± 0.0086 | 0.306 ± 0.0019 | 0.218 ± 0.0053 | 0.215 |
| C4-C4 | 4.8 | 0.271 ± 0.0056 | 0.689 ± 0.0049 | 0.204 ± 0.0026 | 0.051 |
| C1-Nd | 4.0 | 0.297 ± 0.0050 | 0.458 ± 0.0045 | 0.214 ± 0.0014 | 1.486 |
| C1-C2 | 6.7 | 0.322 ± 0.0098 | 0.305 ± 0.0022 | 0.213 ± 0.0035 | 0.174 |
| C2-C2 | 6.6 | 0.342 ± 0.0042 | 0.305 ± 0.0017 | 0.219 ± 0.0017 | 0.154 |
| C3-C3 | 6.0 | 0.307 ± 0.0062 | 0.476 ± 0.0034 | 0.210 ± 0.0022 | 0.154 |
| C1-C4 | 5.8 | 0.317 ± 0.0074 | 0.484 ± 0.0028 | 0.216 ± 0.0030 | 0.173 |
| C3-C4 | 5.4 | 0.295 ± 0.0117 | 0.590 ± 0.0050 | 0.211 ± 0.0028 | 0.151 |
| C2-N0 | 4.3 | 0.300 ± 0.0059 | 0.542 ± 0.0032 | 0.213 ± 0.0015 | 0.072 |
| C3-N0 | 4.0 | 0.290 ± 0.0064 | 0.628 ± 0.0034 | 0.209 ± 0.0020 | 0.051 |
| C2-Nd | 3.9 | 0.288 ± 0.0110 | 0.453 ± 0.0041 | 0.212 ± 0.0040 | 1.527 |
| C2-Nda | 3.9 | 0.310 ± 0.0050 | 0.446 ± 0.0031 | 0.218 ± 0.0017 | 1.404 |
| C3-Na | 3.6 | 0.289 ± 0.0057 | 0.523 ± 0.0033 | 0.212 ± 0.0021 | 1.445 |
| C3-Nda | 3.6 | 0.299 ± 0.0123 | 0.500 ± 0.0026 | 0.213 ± 0.0025 | 1.486 |
| C2-P1 | 2.8 | 0.295 ± 0.0074 | 0.479 ± 0.0043 | 0.213 ± 0.0020 | 1.425 |
| C5-N0 | 2.7 | 0.293 ± 0.0044 | 0.624 ± 0.0028 | 0.212 ± 0.0019 | 1.158 |
| N0-N0 | 2.0 | 0.281 ± 0.0037 | 0.597 ± 0.0044 | 0.204 ± 0.0015 | 1.610 |
| C1-P3 | 1.3 | 0.298 ± 0.0054 | 0.491 ± 0.0032 | 0.212 ± 0.0018 | 1.548 |
| C1-P4 | 1.2 | 0.299 ± 0.0083 | 0.485 ± 0.0027 | 0.211 ± 0.0027 | 1.527 |

### $\Delta G_{\mathrm{Ol}\rightarrow\mathrm{W}}$ for Martini dimers

Figure S3 shows a two dimensional matrix of the octanol/water partition free energy, $\Delta G_{\mathrm{Ol}\rightarrow\mathrm{W}}$, for all combinations of Martini dimers [2].

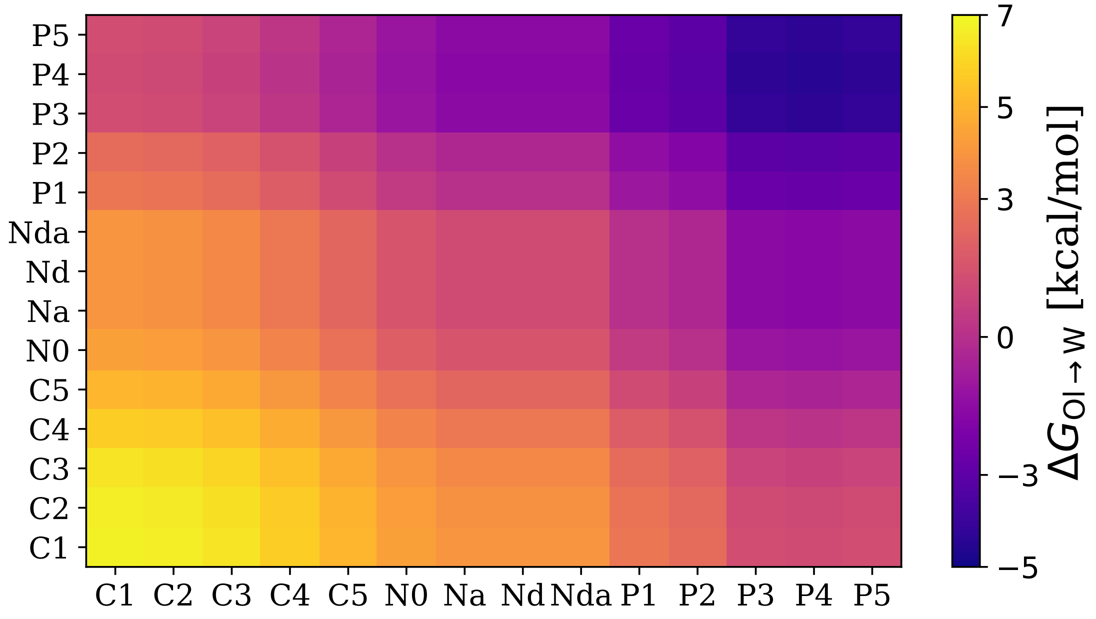


Figure S3: Two dimensional matrix showing $\Delta G_{\mathrm{Ol}\rightarrow\mathrm{W}}$ for all combinations of Martini dimers [2]. Horizontal and vertical axes show the bead type combination of each dimer. The grids are symmetrical across the diagonal.

#### PMFs profiles

Figure S4 shows the $z_{\text{min}}$ location for dimers in minimum (min) environment (a) and median (med) environment (b).

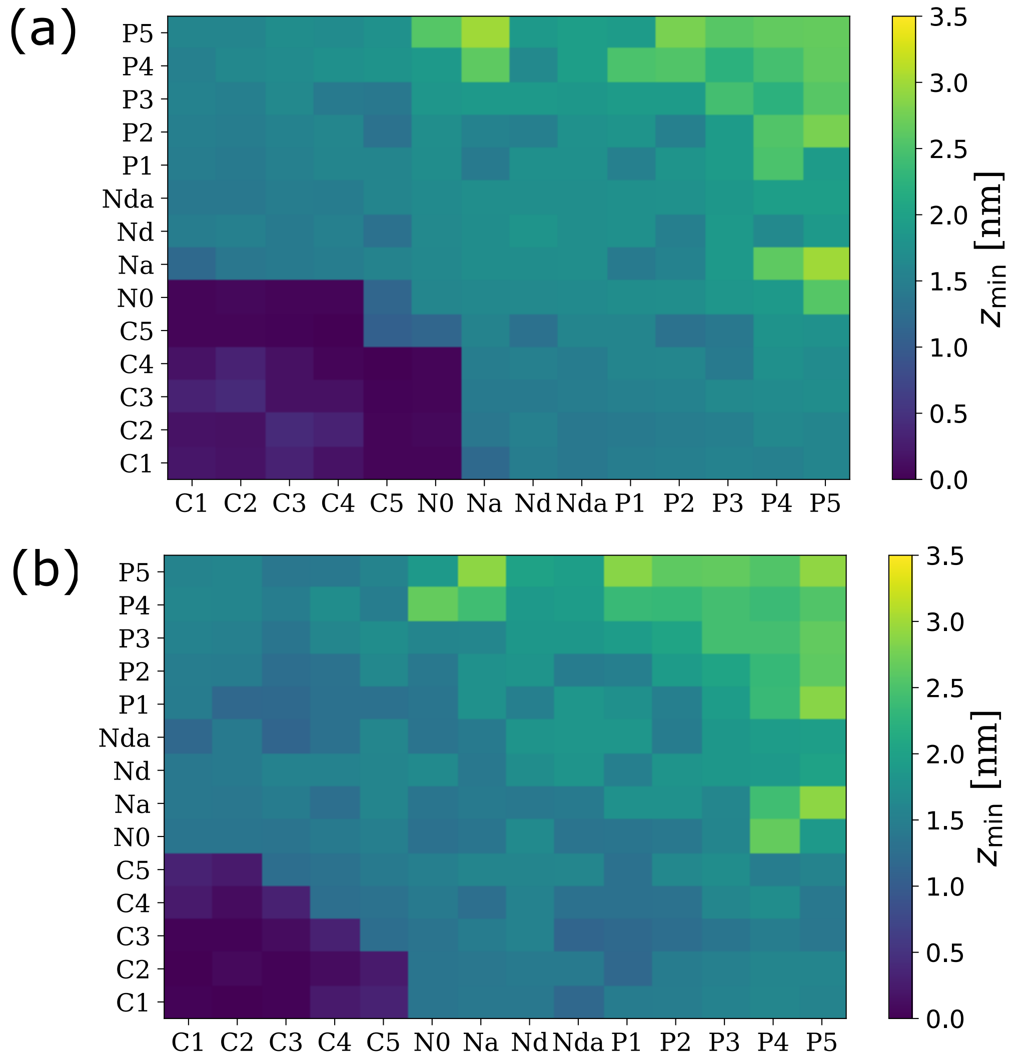


Figure S4: Two dimensional matrices showing $z_{\text{min}}$ location for dimers in the two most favoured environments: minimum (min) environment (a) and median (med) environment (b). Horizontal and vertical axes show the bead type combination of each dimer. The grids are symmetrical across the diagonal.

Figure S5 and Figure S6 show examples of PMF profiles obtained for different Martini dimers in different lipid environments.

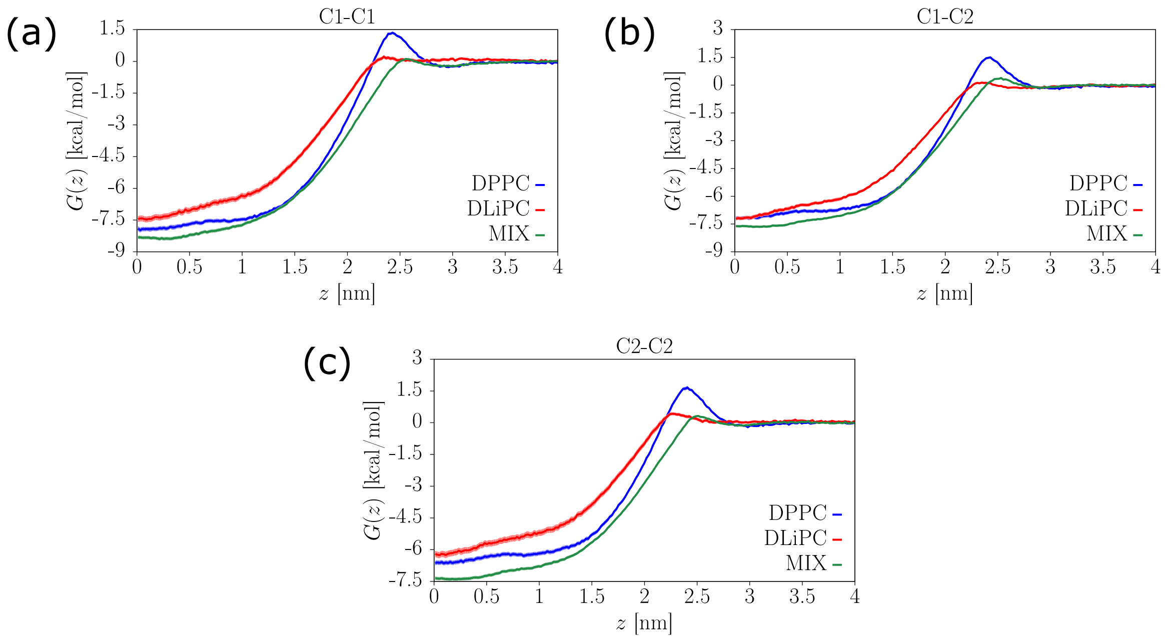


Figure S5: PMF profiles in different lipid environment of dimers falling into the red-square in Figure 4. (a) C1-C1, (b) C1-C2 and (c) C2-C2.

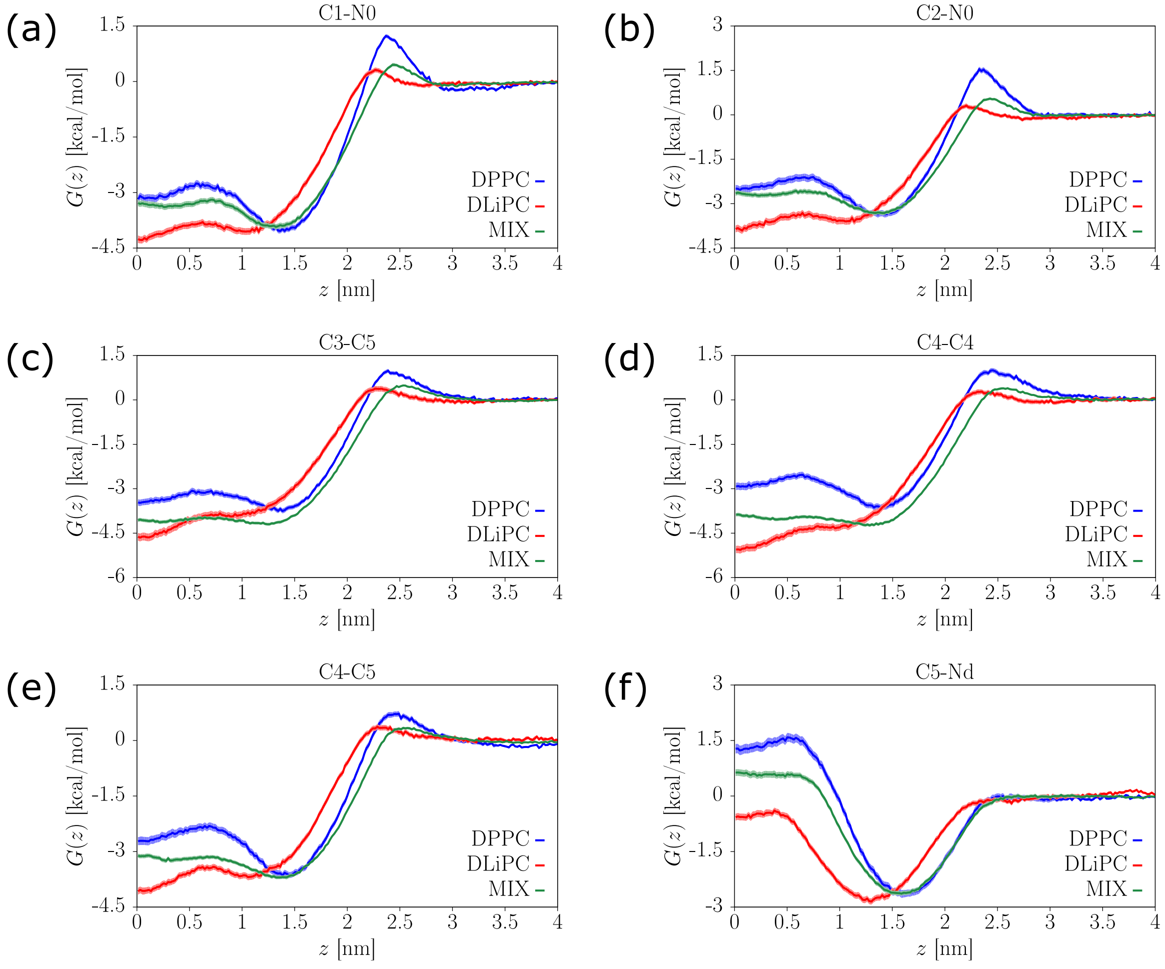


Figure S6: PMF profiles in different lipid environment of dimers falling into the blue-diagonal in Figure 4. (a) C1-N0, (b) C2-N0, (c) C3-C5, (d) C4-C4, (e) C4-C5 and comparison with a dimer not belonging to the blue-diagonal (f) C5-Nd.

#### Potentials of mean force in different lipid environments: effect of cholesterol

Figure S7 shows the $\Delta G_{\mathrm{W}\to\mathrm{z_m}}$ obtained for different Martini dimers in the DPPC-DLiPC-CHOL systems at different cholesterol concentrations.

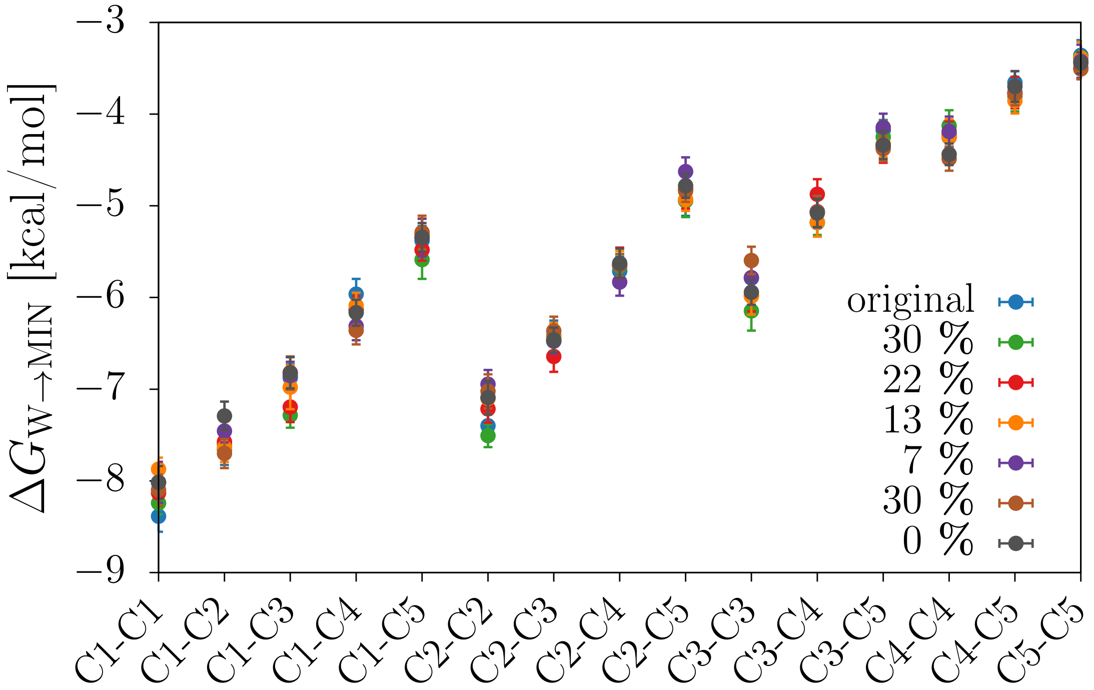


Figure S7: $\Delta G_{\mathrm{W}\to\mathrm{z_m}}$ obtained for the original system (41 % DPPC, 29 % DLiPC and 30 % CHOL) and for ternary membranes with equal ratios of DPPC and DLiPC and different cholesterol concentration (0, 3, 7, 13, 22 and 30 mol%).

Figure S8 and Figure S9 show matrices for the $\Delta\Delta G$ and $z_{\text{min}}$ location obtained for different Martini dimers in the DPPC-DLiPC-CHOL systems at different cholesterol concentrations.

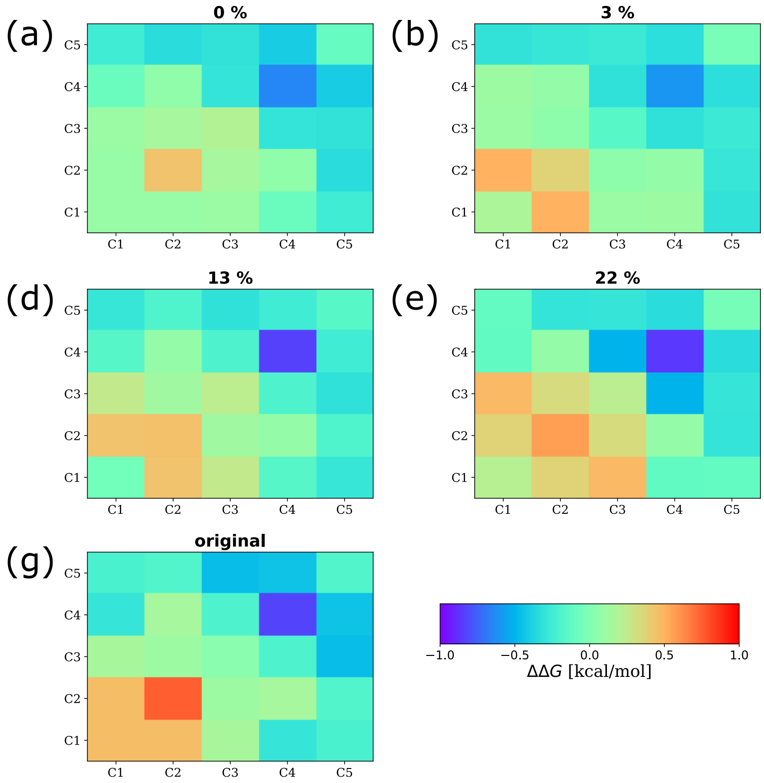


Figure S8: Two dimensional matrices showing $\Delta\Delta G$ in the ternary membranes with equal ratios of DPPC and DLiPC and different cholesterol concentration (0, 3, 7, 13, 22 and 30 mol%) in comparison to the original system (41 % DPPC, 29 % DLiPC and 30 % CHOL). The colour scale is the same in all figures.

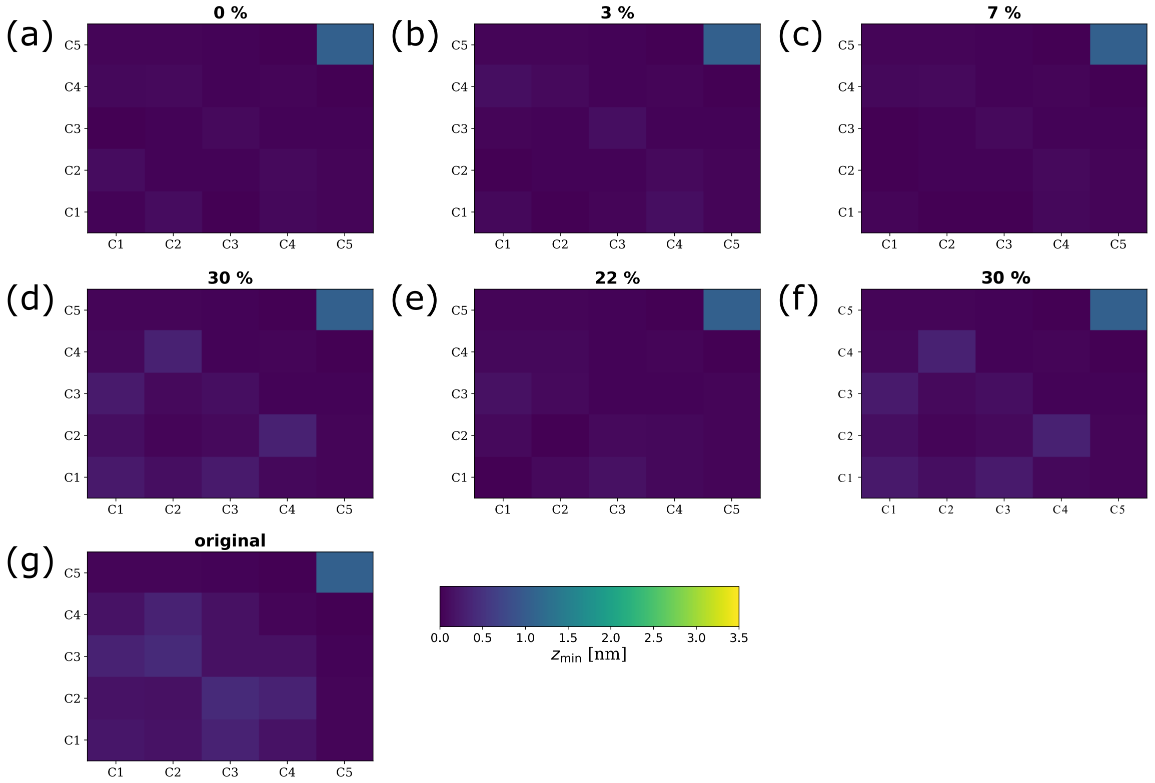


Figure S9: Two dimensional matrices showing $z_{\text{min}}$ location for dimers in the minimum (min) environment in membranes at different cholesterol concentrations. Horizontal and vertical axes show the bead type combination of each dimer. The grids are symmetrical across the diagonal.

### Mean-Field Theory

(*A Note about notations:* Allender and Schick have used $T_{\text{C}}$ as the critical mixing temperature while we have used $T_{\text{mix}}$. We are using $T_{\text{C}}$ in this SI to be consistent with their notations.)

In their paper, Allender and Schick [1] modeled the lipid membrane as a polymer blend composed of three linear homopolymers: an unsaturated lipid (A), a saturated lipid (B), and the solute (S). To find the thermodynamic properties of the system at equilibrium they then used a Flory-Huggins type Helmholtz free energy (Eq. (2) of their paper):

$$F(T, n_{\text{A}}, n_{\text{B}}, n_{\text{S}}) = V_{\text{AB}} n_{\text{A}} N_{\text{A}} \frac{n_{\text{B}} N_{\text{B}} v_0}{\Omega} + V_{\text{AS}} n_{\text{A}} N_{\text{A}} \frac{n_{\text{S}} N_{\text{S}} v_0}{\Omega} + V_{\text{BS}} n_{\text{B}} N_{\text{B}} \frac{n_{\text{S}} N_{\text{S}} v_0}{\Omega} + k_{\text{B}} T \sum_{i=A,B,S} n_i \ln\left(\frac{n_i N_i v_0}{\Omega}\right)$$

where $v_0$ is the monomer volume (assumed to be the same for all lipids and the solute), $n_i$ and $N_i$ (with $i = \text{A}, \text{B}$ or S) are the number of molecules and degrees of polymerization of unsaturated (A), saturated (B), and solute (S) molecules, respectively, $\Omega = v_0(n_{\text{A}} N_{\text{A}} + n_{\text{B}} N_{\text{B}} + n_{\text{S}} N_{\text{S}})$ is the total volume of the system and $k_{\text{B}}$ is the Boltzmann's constant. The interaction energy $V_{\text{AB}}$ is defined as

$$V_{\text{AB}} = \tilde{V}_{\text{AB}} - \frac{(\tilde{V}_{\text{AA}} + \tilde{V}_{\text{BB}})}{2},$$

in which $\tilde{V}_{\mathrm{AA}}$ and $\tilde{V}_{\mathrm{BB}}$ are interactions between pairs of A monomers and B monomers, respectively. $\tilde{V}_{\mathrm{AB}}$ is the interaction energy between AB pairs. Note that $V_{\mathrm{AB}}$ is proportional to the standard Flory $\chi$ parameter.

Then equating the chemical potentials of similar type of molecules on the critical line across the coexistence surface of Ld-rich phase and Lo-rich phase, they found the critical partitioning ($M_{\mathrm{C}}$) and the change in critical mixing temperature ($\Delta T$) as a function of the solute volume fraction $\Phi_{\mathrm{S}}$, which in the limit of dilute solute concentration become (see Eq. (17) and Eq. (21) of Allender and Schick paper, respectively)

$$M_{\mathrm{C}} = -\frac{N_{\mathrm{S}} V_{\mathrm{AB}}}{k_{\mathrm{B}} T_{\mathrm{C}}(0)} (\delta\nu + \delta r)\Phi_{\mathrm{S}} + O(\Phi_{\mathrm{S}}^2), \tag{S1}$$

$$\Delta T \equiv \frac{T_{\mathrm{C}}(\Phi_{\mathrm{S}}) - T_{\mathrm{C}}(0)}{T_{\mathrm{C}}(0)} = \Phi_{\mathrm{S}} \left[ -1 + \frac{N_{\mathrm{S}} V_{\mathrm{AB}}}{2 k_{\mathrm{B}} T_{\mathrm{C}}(0)} (\delta\nu + \delta r)^2 \right], \tag{S2}$$

in which $T_{\mathrm{C}}(0)$ is the critical temperature of the bilayer without any solute.

We evaluate $(\delta\nu + \delta r)$ from Eq. (S1) and use it to rewrite Eq. (S2) as

$$\Delta T = \Phi_{\mathrm{S}} \left[ -1 + \alpha \frac{M_{\mathrm{C}}^2}{\Phi_{\mathrm{S}}^2} \right], \text{ where } \alpha \equiv \frac{k_{\mathrm{B}} T_{\mathrm{C}}(0)}{2 N_{\mathrm{S}} V_{\mathrm{AB}}}. \tag{S3}$$

Slightly changing this equation (using $T_{\mathrm{mix}}$ as the critical mixing temperature instead of $T_{\mathrm{C}}$, as mentioned in the beginning of this section and redefining $\Delta T$ as temperature difference rather than a dimensionless ratio), we use the following form of it in the main text (Eq. (4)):

$$\frac{\Delta T_{\mathrm{mix}}}{T_{\mathrm{mix}}(0)} \equiv \frac{T_{\mathrm{mix}}(\Phi_{\mathrm{S}}) - T_{\mathrm{mix}}(0)}{T_{\mathrm{mix}}(0)} = \Phi_{\mathrm{S}} \left[ -1 + \alpha \frac{M_{\mathrm{C}}^2}{\Phi_{\mathrm{S}}^2} \right].$$

#### Comparison between MD simulations and predictions from mean-field theory

Recall that $M_{\mathrm{C}}$ was defined as:

$$M_{\mathrm{C}} = \left. \frac{\Phi_{\mathrm{S}}^{\mathrm{II}} - \Phi_{\mathrm{S}}^{\mathrm{I}}}{\Phi_{\mathrm{A}}^{\mathrm{II}} - \Phi_{\mathrm{A}}^{\mathrm{I}}} \right|_{\mathrm{C}} \tag{S4}$$

where $\Phi_{\mathrm{S}}^{\mathrm{II}}$, $\Phi_{\mathrm{A}}^{\mathrm{II}}$, $\Phi_{\mathrm{S}}^{\mathrm{I}}$ and $\Phi_{\mathrm{A}}^{\mathrm{I}}$ are the critical volume fractions of solute and lipid A in phase II (i.e. the A-rich Ld phase) and in phase I (i.e. the B-rich Lo phase), respectively. To connect mean-field and MD simulations, volume fractions are approximated with the ratio of contacts, $C_{i-j}$, as follows:

$$\Phi_{\mathrm{S}}^{\mathrm{I}} = \frac{C_{\text{S-B}}}{C_{\text{A-B}} + C_{\text{B-B}} + C_{\text{S-B}}}, \tag{S5}$$

$$\Phi_{\mathrm{S}}^{\mathrm{II}} = \frac{C_{\text{S-A}}}{C_{\text{A-A}} + C_{\text{B-A}} + C_{\text{S-A}}}, \tag{S6}$$

$$\Phi_{\mathrm{A}}^{\mathrm{I}} = \frac{C_{\text{A-B}}}{C_{\text{A-B}} + C_{\text{B-B}} + C_{\text{S-B}}}, \tag{S7}$$

$$\Phi_{\mathrm{A}}^{\mathrm{II}} = \frac{C_{\text{A-A}}}{C_{\text{A-A}} + C_{\text{B-A}} + C_{\text{S-A}}} \tag{S8}$$

where A is DLiPC, B is DPPC and S is the dimer. It should be noted that $C_{\text{A-B}}$ and $C_{\text{B-A}}$ are equivalent. Each contact, $C_{i-j}$, is measured from simulations following the same protocol described in Contact fraction analysis, obtaining the volume fractions in Table S3 from which $M_{\mathrm{C}}$ values can be calculated as shown in Figure S10.

Table S3: Volume fractions, $\Phi$, as defined in Eq. S5- S8 measured from MD simulations of the system DPPC:DLiPC:CHOL in the presence of dimers, critical partitioning, $M_{\mathrm{C}}$, calculated from Eq. S4 and $\Delta T_{\mathrm{mix}}$ obtained from Eq. (4) of the main text by fixing the adjustable parameter $\alpha = 0.003$. A is DLiPC, B is DPPC, S is the dimer, I is the B-rich Lo phase and II is the A-rich Ld phase.

| Dimer | $\Phi_{\mathrm{A}}^{\mathrm{I}}$ | $\Phi_{\mathrm{A}}^{\mathrm{II}}$ | $\Phi_{\mathrm{S}}^{\mathrm{I}}$ | $\Phi_{\mathrm{S}}^{\mathrm{II}}$ | $M_{\mathrm{C}}$ | $\Delta T_{\mathrm{mix}}$ |
|---|---|---|---|---|---|---|
| C1-C1 | 0.135 | 0.471 | 0.332 | 0.275 | -0.170 | -0.0162 |
| C1-C2 | 0.126 | 0.494 | 0.334 | 0.271 | -0.171 | -0.0162 |
| C1-C4 | 0.135 | 0.421 | 0.281 | 0.383 | 0.357 | 0.00060 |
| C2-C2 | 0.134 | 0.480 | 0.333 | 0.271 | -0.179 | -0.0159 |
| C2-N0 | 0.133 | 0.415 | 0.257 | 0.407 | 0.532 | 0.0102 |
| C3-C3 | 0.130 | 0.435 | 0.279 | 0.373 | 0.308 | -0.0094 |
| C3-C4 | 0.132 | 0.402 | 0.237 | 0.431 | 0.719 | 0.0345 |
| C3-N0 | 0.134 | 0.395 | 0.218 | 0.444 | 0.866 | 0.0587 |
| C4-C4 | 0.129 | 0.382 | 0.195 | 0.476 | 1.111 | 0.1091 |

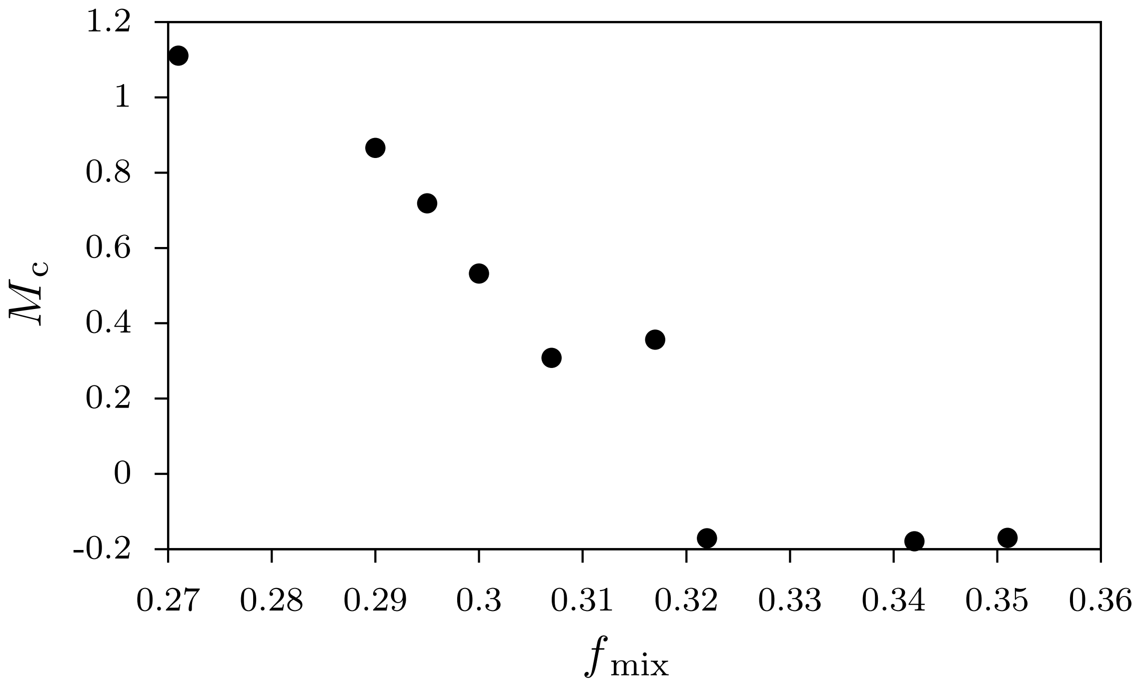


Figure S10: Critical partitioning, $M_{\mathrm{c}}$, as a function of the system contact fraction, $f_{\mathrm{mix}}$.

[2] T. Bereau and K. Kremer. Automated parametrization of the coarse-grained martini force field for small organic molecules. 11(6):2783–2791.
